# Durable reconstitution of sinonasal epithelium by transplant of CFTR gene corrected airway stem cells

**DOI:** 10.1101/2025.01.24.634776

**Authors:** Dawn T. Bravo, Sriram Vaidyanathan, Jeannette Baker, Vrishti Sinha, Esmond Tsai, Pooya Roozdar, William W. Kong, Patrick J Atkinson, Zara M. Patel, Peter H. Hwang, Vidya K. Rao, Robert S. Negrin, Jeffrey J. Wine, Carlos Milla, Zachary M. Sellers, Tushar J. Desai, Matthew H. Porteus, Jayakar V. Nayak

## Abstract

Modulator agents that restore cystic fibrosis transmembrane conductance regulator (CFTR) function have revolutionized outcomes in cystic fibrosis, an incurable multisystem disease. Barriers exist to modulator use, making local CFTR gene and cell therapies attractive, especially in the respiratory tract. We used CRISPR to gene-correct CFTR in upper airway basal stem cells (UABCs) and show durable local engraftment into recipient murine respiratory epithelium. Interestingly, the human cells recapitulate the *in vivo* organization and differentiation of human sinus epithelium, with little expansion or contraction of the engrafted population over time, while retaining expression of the CFTR transgene. Our results indicate that human airway stem cell transplantation with locoregional restoration of CFTR function is a feasible approach for treating CF and potentially other diseases of the respiratory tract.

## Introduction

Cystic fibrosis (CF) is an incurable, life-limiting, multisystem disease caused by variants in the cystic fibrosis transmembrane conductance regulator (*CFTR*) gene (*1*). Treatment of CF has been revolutionized by small molecule drugs that augment CFTR function, thereby improving anion transport on mucosal surfaces. While attention to date has primarily focused on pulmonary disease (*2*), CF-related chronic rhinosinusitis (CF-CRS) is a major source of morbidity in nearly all people with CF (*3–5*). CF-CRS is characterized by impaired mucociliary clearance, leading to chronic sinonasal tissue inflammation and bacterial overgrowth of drug-resistant pathogens (*6*–*9*). Overlapping microbial isolates between the sinuses and lungs suggest that purulent discharge from the upper airway can cause chronic contamination of the pulmonary tree (*10–12*). Active treatments for CF-CRS include topical rinses, antibiotics, and repeated sinus surgeries to clear inflammation and reduce bacterial burden (*13*). Ameliorating CF-CRS is a pressing concern that may have direct implications on pulmonary disease progression in all people with CF (*14*). The transformational impact on CF seen with the advent of modulator therapy indicates that increased expression of CFTR can greatly reduce disease burden, while also highlighting the need for novel solutions to address barriers to therapy including i) modulator non-responsiveness based on genotype, especially in underserved populations (15); (ii) lifelong medication need and medication costs (16); and iii) notable rates of toxicities and intolerance (17, 18).

Collectively, these observations demonstrate a pressing need to develop a stem cell or gene therapy to durably augment CFTR function in the airway epithelium. Past gene therapy approaches to drive airway CFTR expression have been unsuccessful (*19–22*), and the *in vivo* delivery of Cas9 poses concerns about anti-Cas9 immune responses (*23–25*). Considering these limitations, we have advanced the innovative concept of *ex vivo CFTR* gene correction in primary upper airway basal stem cells (UABCs) isolated from CF donors, followed by staged, autologous, re-transplantation into upper airways. Given the substantial hurdles inherent to delivery and engraftment of airway stem cells into the lungs of CF patients, efforts were steered towards treatment of CF-CRS, a relevant and approachable respiratory target site in CF. We have previously demonstrated efficient *CFTR* gene correction of the common F508del mutation in primary CF UABCs using CRISPR/Cas9 (*26*), and restoration of CFTR function in human UABCs to levels seen in non-CF controls using full length CFTR cDNA (*27*). Importantly, gene-corrected UABCs properly differentiated into ciliated and mucus-producing airway epithelial cells. We now report our progress with *in vivo* transplantation and stable engraftment of primary UABCs derived from transgenic reporter mice, and gene-corrected primary UABCs from human CF/non-CF patients into recipient immunocompromised mice. These advances serve as a pre-clinical springboard towards first-in-human stem cell trials to treat a respiratory disease, specifically CF-CRS.

## Results

### Transplantation and engraftment of syngeneic murine UABCs to the injured upper airway of mice to reconstitute the normal epithelium

A workflow for *ex vivo* harvest and *in vivo* transplantation of murine UABCs is illustrated (Figure 1A). Transgenic mice constitutively co-expressing GFP and luciferase (GFP-Luc) were first exploited to properly track UABCs (*28*, *29*). Nasal epithelial disruption of recipient immunocompromised NSG mice was performed using various injuries: chemical with sulfur dioxide (SO_2_) gas (Fig. 1B and fig. S1, S2), mechanical with nasal brush, (NB) placement (Fig. 1C and fig. S3) (*30*, *31*), or detergent based with 2% polidocanol (PDOC) (Fig. 1D) (*32*), GFP-Luc UABCs encapsulated in Matrigel^TM^ were instilled unilaterally into the mouse nostril. UABCs engrafted into the nasal airways within 2 weeks based on prominent bioluminescence signal, which was maintained over 6 (Fig. 1B, C) and 9 months (Fig. 1D) thus demonstrating durable engraftment. Radiance measurements (photons/sec) recordings confirmed highly significant, durable engraftments for all 3 preconditioning treatments (p=0.0001), (Fig. 1E, fig. S2 A&B and fig. S3A). Given ease and reproducibility of nasal epithelial injury using 2% PDOC, all further recipient mouse nasal preconditioning was performed using this topical sclerosing agent.

**Fig. 1:**
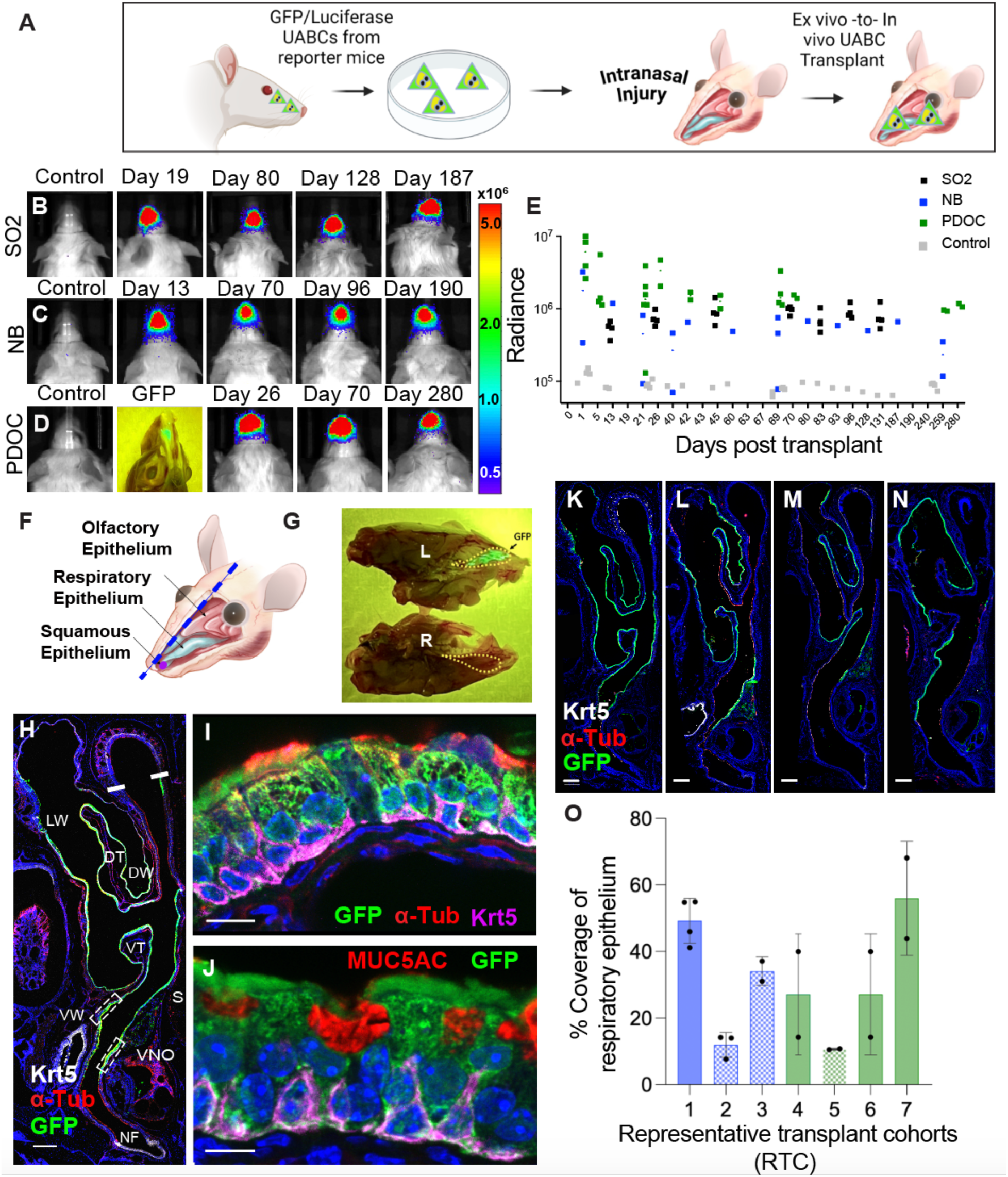
Durable engraftment and epithelial reconstitution following *ex vivo* to *in vivo* transplantation of murine UABCs. (**A**) A workflow for *ex vivo* harvest and *in vivo* transplantation of murine UABCs is illustrated. (**B-D**) Bioluminescent images (BLI) demonstrate stable engraftment of UABCs derived from transgenic reporter mice constitutively expressing GFP and luciferase (GFP-Luc) to the upper airways of NSG mice following preconditioning by (B) chemical (SO_2_ gas), (C) mechanical nasal brush (NB), and (D) sclerosing agent polidocanol (PDOC). (**E**) Graphical representation of BLIs exhibit highly significant, stable and durable engraftment for over 6-9 months (p <0.0001). (**F-G**) Transplanted GFP-Luc UABCs show isolated, controlled engraftment to the respiratory epithelium of recipient NSG mice as seen by Stereo Fluorescent Microscopy (SFM) showing GFP positive cells (G, left (L)). SFM also excites GFP signal through the decalcified skull (C). (**H-J**) Immunofluorescent microscopy demonstrates precise linear engraftment of GFP+ cells solely along the upper airway epithelial basement membrane of NSG transplanted mice (H). Magnified view of (H) demonstrates epithelial reconstitution from transplanted GFP-Luc UABCs (I&J) co-expressing the basal cell protein cytokeratin 5+ (I, Krt5, magenta), as well as differentiated, luminal ciliated cells (I, α-tubulin, red). The magnified view of dashed box adjacent to the vomeronasal organ (VNO, H), shows GFP+ progeny cells expressing mucin polysaccharide (J, MUC5AC, red) found in differentiated secretory cells. Because of plane of focus, Krt5 expression is not equivalent in all sections. (**K-N**) Representative transplant cohorts (RTCs) display an expected range of localized engraftment of transplanted GFP reporter cells with the central airways. (**O**) The percent coverage of respiratory epithelium was determined to be ∼50% in some RTCs. Engraftment was equivalent between male (blue) or female (green) mice (O). More efficient engraftments were obtained using fewer than 1×10^6^ UABCs (N, patterned bars). Anatomical regions of the nasal cavity/upper airways: Ventral Wall (VW), septum (S), ventral turbinate (VT), dorsal turbinates (DT), lateral wall (LW) and dorsal wall (DW). Scale bars: 100 µm (H, K-N) 10 µm (I&J).

Sagittal sections of skulls from transplanted recipient mice (injured using 2% PDOC) demonstrates intranasal GFP signal confined to the unilateral respiratory nasal cavity using stereo fluorescent microscopy (SFM) (Fig. 1F, G, yellow dashed line, also Fig. 1D). At low resolution, cross sections of the recipient mouse nasal cavity show a widespread linear staining pattern for GFP and cytokeratin 5 (Krt5) at the outer perimeter of the nasal septum, turbinate and side wall structures (Fig. 1H). At higher magnification (Fig. 1H, dashed boxes) instilled GFP-Luc UABC basal stem cells show engraftment to the basement membrane layer of the respiratory epithelium (Fig. 1I, J). Progeny differentiated GFP^+^ ciliated cells co-expressing acetylated tubulin (α-tub, Fig. 1I) and secretory cells expressing mucin 5AC (MUC5AC, Fig. 1J) also arose to reconstitute the native epithelial architecture. Similar patterns of UABC stem cell engraftment to the basement membrane were noted after NB or SO_2_ treatment (fig. S2C, S3 B&C). Most reporter UABCs were seen to engraft to central unilateral nasal airway structures following transplant (Figure 1H, K-N). Approximately 150 serial sections from the central nasal cavity were obtained from 7 representative transplant cohorts (RTC) to gain comprehensive insight of the degree of engraftment. Mean engraftment was approximately 30% (29.9% ± 9.9%, Fig. 1O), with greater than 50% epithelial coverage quantified for several mice in 2 cohorts (Fig 1O, 1&7 RTC, 52.3%+/- 4.8%). No difference in engraftment was seen between recipient mice by sex (Fig. 1O, blue vs green bars, (1.09% +/- 0.64%), and there was more efficient engraftment via installation of approximately 1×10^6^ starting cells (Fig 1O, pattern bars).

### Fibrinogen and thrombin support the proliferation of murine UABCs, and effective engraftment *in vivo*

Following successful stable engraftment of UABCs to sites of upper respiratory injury using Matrigel™, we next identified a clinically applicable scaffold for human basal stem cell delivery. A range of matrix biomaterials were assayed to promote human UABC stem cell replication and survival (Fig. 2A, Supplementary Table 1, (*26*, *33–36*)). Human UABCs proliferated most actively when cultured in either fibrinogen or laminin (BioSilk^TM^, Fig. 2A) while still retaining the expression of stem cell biomarkers Krt5 and nuclear transcription factor p63 (Fig. 2B-E). Varied concentrations of fibrinogen (Fig. 2F) and thrombin (Fig. 2G) were optimized for maintenance of murine UABC cell proliferation. Subsequent transplant experiments were performed with gels containing admixed fibrinogen with thrombin matrix.

**Fig. 2:**
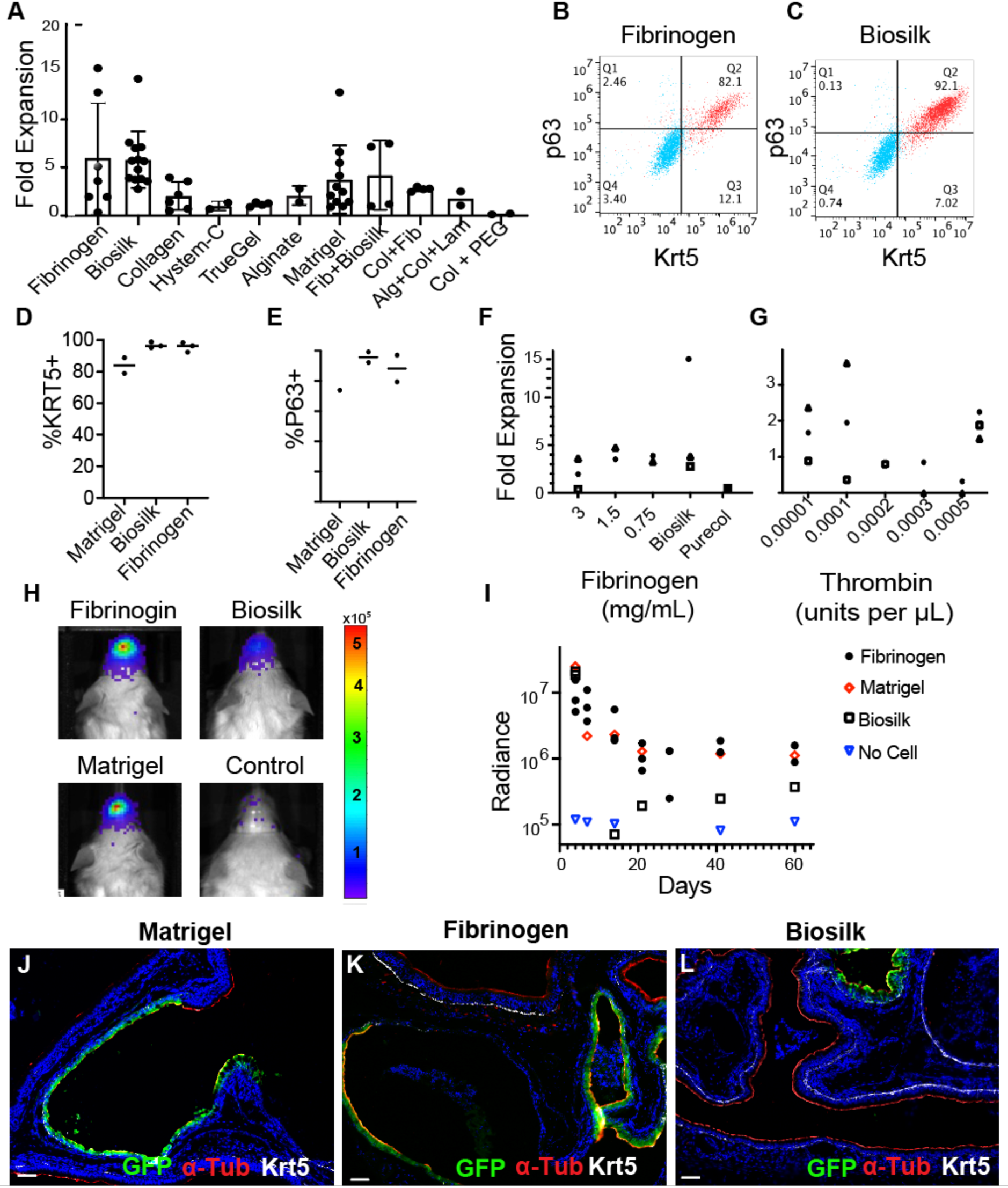
Fibrinogen and thrombin support optimal engraftment of UABCs. (**A**) The expansion of human UABCs was investigated in a variety of biomatrices for optimal proliferation. UABCs proliferated in human fibrinogen and recombinant Laminin gel (Biosilk^TM^) at rates comparable to Matrigel^TM^. (**B-E**) UABCs cultured in fibrinogen and Biosilk^TM^ retained the expression of KRT5 and p63 stem cell biomarkers. Representative images show KRT5 and p63 expression relative to isotype controls (B&C). (**D, E**) Data from additional donors. (**F**) Fibrinogen gels between concentrations of 0.75 mg/mL – 3 mg/mL were evaluated. The proliferation of UABCs was not significantly different between different concentrations. (**G**) The amount of thrombin impacts the rigidity of the gel and the gelation time. We evaluated the impact of 0.1 – 5 * 10^-4^ units of thrombin per μL (0.004 – 0.02 units of thrombin per 40 uL dome). Concentrations > 2 * 10^-4^ units of thrombin per μL reduced the proliferation of UABCs. (**H, I**) Mouse UABCs were encapsulated in fibrinogen, Biosilk or Matrigel^TM^ and transplanted after injury using 2% polidocanol (PDOC). Stable engraftment was observed 60 days post-transplant. Matrigel^TM^ and fibrinogen demonstrate similar engraftment levels (I). Bioluminescence was measured in mice transplanted with murine UABCs using the different biomaterials. (**J-L**) Representative immunofluorescence images showing engraftment of UABCs GFP-Luc cells transplanted in Matrigel^TM^ (J) fibrinogen (K), or Biosilk (L). Similar degrees of healthy stem cell engraftment and epithelial architecture was conferred in UABCs transplanted in Matrigel^TM^ and fibrinogen compared to Biosilk. Scale bars: 25µm.

When comparing fibrinogen vs. Biosilk^TM^ to facilitate *in vivo* engraftment in NSG mice after PDOC injury, GFP-Luc UABCs encapsulated in fibrinogen showed similar radiance signal to Matrigel^TM^ at 60 days post-transplant, in contrast to upper airway stem cells transplanted with Biosilk^TM^ (Fig. 2H-I). At the cellular level, murine GFP-Luc UABCs transplanted in either fibrinogen or Matrigel^TM^ generated well-differentiated nasal epithelium with robust Krt5 and α-tubulin expression in GFP+ cells, while delivery in Biosilk^TM^ appeared to generate a rudimentary epithelium (Fig. 2J-L). Since fibrinogen is a human protein, we used fibrinogen in subsequent experiments.

### Engineering of human UABCs to permit *in vivo* detection

Schematic workflow for *ex vivo* surgical retrieval and transplantation of human CF/non-CF UABCs is illustrated (Figure 3A). To track human stem cells following xenotransplant, we first engineered primary human UABCs using CRISPR/Cas9 and AAV to constitutively express both GFP and Nanoluc^TM^ (GFP-NanoLuc) from the *HBB* locus (Fig. 3B), a ‘safe harbor’ in basal stem cells given absence of HBB gene expression in the airway epithelium. GFP-NanoLuc-expressing human UABCs were enriched using fluorescence activated cell sorting (FACS) to ∼80% GFP+ cells (Fig. 3B, C), admixed with fibrinogen/thrombin, and transplanted into the upper airways of NSG mice following PDOC treatment. GFP-NanoLuc human UABCs provided stable bioluminescence signals for 4+ months (Fig. 3D). Radiance measurements were significantly higher in the upper airways of engrafted UABC recipient mice compared to controls (p=0.018) (Fig. 3D, E, fig. S4).

**Fig. 3:**
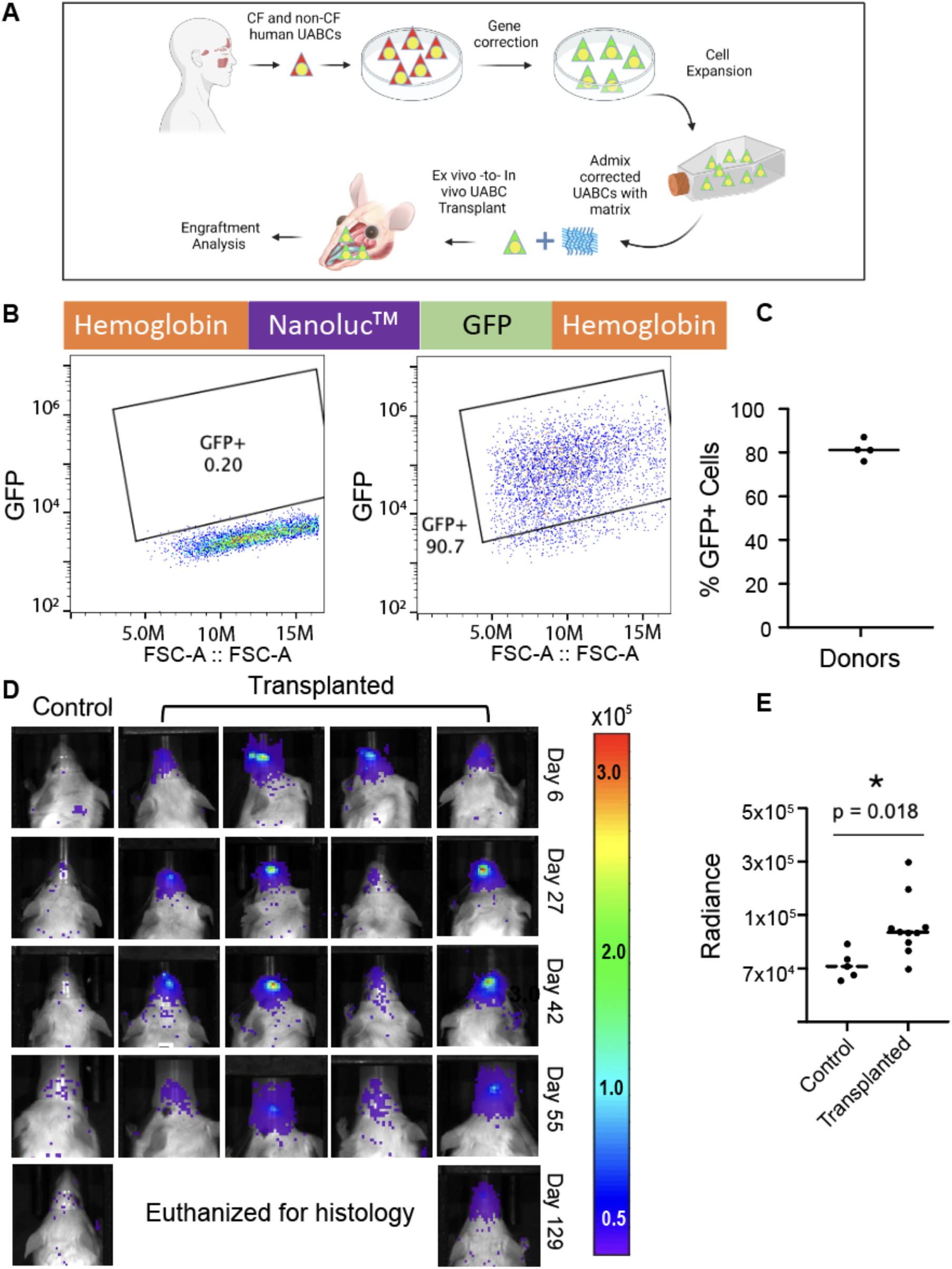
Transplantation of human UABCs engineered to express GFP and Luciferase. (**A**) Schematic illustration for *ex vivo* surgical retrieval and *in vivo* transplantation of human CF/non-CF UABCs. (**B**) Human UABCs were edited to express Nanoluc^TM^ luciferase and GFP from the beta-hemoglobin (HBB) locus. Representative FACS plots demonstrate GFP expression in ∼90% of the enriched cells (**C**) UABCs from multiple donors edited to express Nanoluc^TM^ luciferase and GFP, show >80% GFP expression following enrichment. (**D**) Edited UABCs edited to express GFP-Luc stably engraft into the murine upper airway over 4+ months. (**E**) Several recipient mice demonstrated significant *in vivo* engraftment above controls measured via bioluminescence live imaging (p=0.018, asterisk).

### Transplantation and detection of human *CFTR* gene corrected UABCs (gcUABCs)

The stable engraftment of GFP-NanoLuc reporter human UABCs provided a road map for human airway stem cell transplantation into mice. To advance this application to CF, we used a large transgene knock-in system in human UABCs from CF and non-CF donors as previously described (*27*) (Fig. 4A). In this knock-in editing system, the full-length human CFTR cDNA is inserted precisely into the endogenous CFTR start codon along with an expression cassette for truncated CD19 (tCD19) rendered devoid of intracellular physiologic signaling. tCD19+ gene corrected UABCs (gcUABCs) were readily detected on flow cytometry (Fig. 4A), and enriched to 57±4% corrected cells using magnetic activated cell sorting (MACS) (Fig. 4B). To confirm functionality after knock-in and enrichment, sorted gcUABCs were differentiated in air-liquid interface (ALI) cultures to assess for CFTR function. Mean CFTR_inh_-172 sensitive short circuit current was 11±10µA/cm^2^, following gene correction compared to 1±1 µA/cm^2^ in CF samples prior to correction (Fig. 4C, D). CFTR_inh_-172 sensitive short circuit current in non-CF controls was 19±11 µA/cm^2^ (Figure 4D and Supplementary Table 2). Thus, in ALI cultures the measured physiologic correction exceeded the predicted 20% improvement required for rescue of aberrant epithelial physiology in CF, and the measured functional rescue was directly proportional to the gene correction frequency following selection (57±4% at 11 µ A/cm^2^ vs. 58% at 19 µA/cm^2^).

**Fig. 4:**
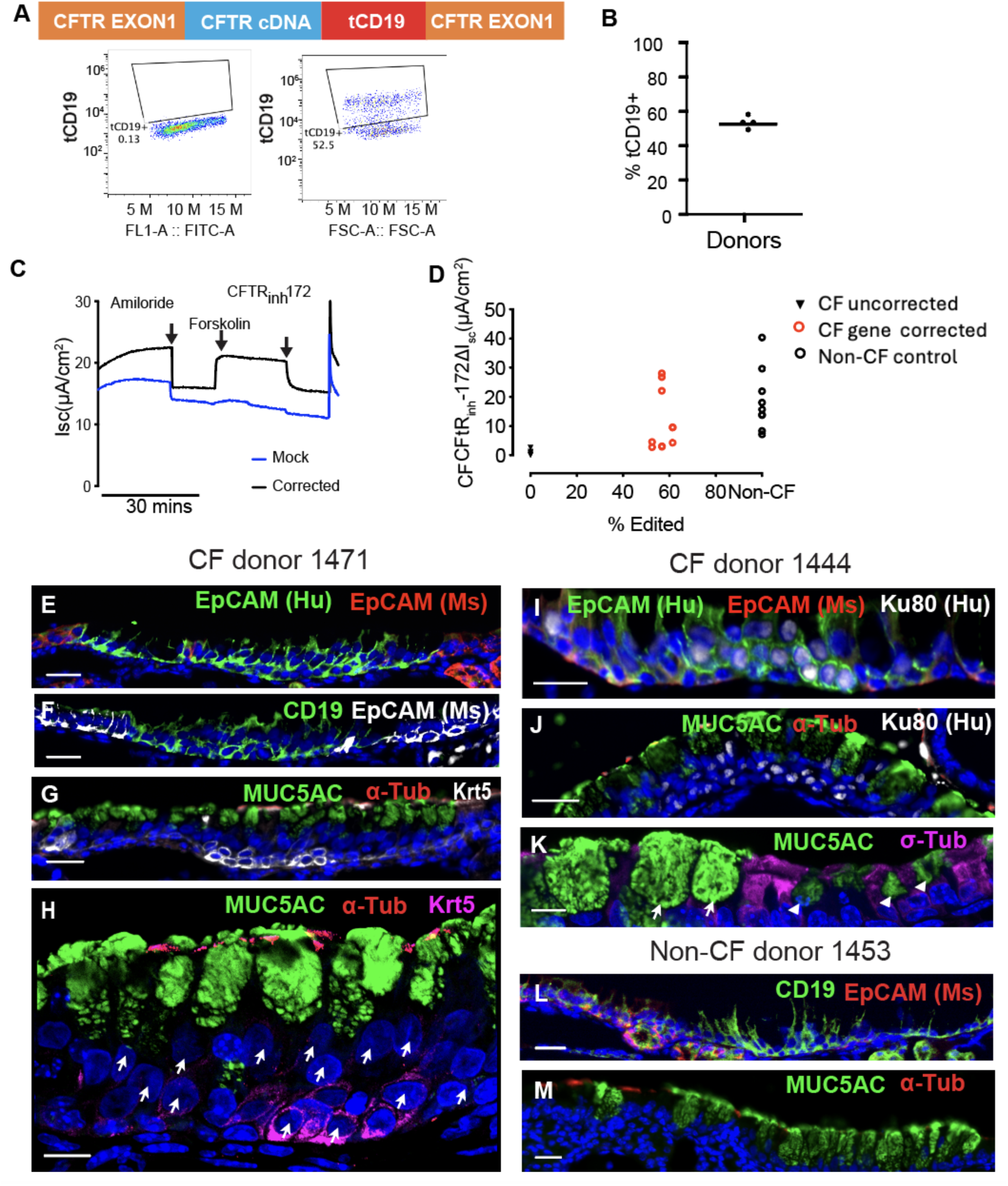
Successful engraftment and airway epithelial reconstitution of *ex vivo* gene corrected human UABCs from CF donors. (**A**) UABCs from donors with and without CF disease were edited to express the full-length CFTR cDNA and truncated CD19 enrichment tag at the endogenous CFTR locus. Representative FACS plots demonstrated that >50% of enriched UABCs were tCD19+ after knock-in compared to isotype controls. (**B**) Magnetic activated cell sorting (MACS) was used to enrich tCD19-expressing UABCs to >50% corrected cells from multiple donors. (**C**) Enriched gcUABCs were grown and differentiated in air liquid interface (ALI) cultures, and representative traces of restored CFTR function measured using Ussing chamber assay (black tracing). (**D**) Cultured gcUABCs demonstrated restored CFTR function as determined by short-circuit currents similar to CFTR function in non-CF control epithelia. (**E-G**) Serial epithelial cross sections from recipient mice transplanted with gcUABCs from CF donor 1471 demonstrate engrafted gcCFTR UABCs. (**E**) Human specific epithelial cell adhesion molecule (EpCAM Hu, green) distinguishes niches engrafted with gcCFTR UABCs while mouse specific EpCAM (EpCAM Ms, red) sharply delineates recipient mouse respiratory epithelium at the niche perimeter. (**F**) Human gcCFTR UABCs in this niche also co-express truncated CD19 (tCD19, green) flanked by mouse epithelium (EpCAM Ms, white). (**G**) Differentiated, secretory (MUC5AC, green) and ciliated cells (α-tubulin, red), also arise from gcCFTR UABCs within this space, with Krt5 (white) expressed along the niche basement membrane. Tiers of Krt5+ UABCs are characteristic of normal human upper airway tissues. (**H**) High resolution image of (G) demonstrating human nuclei (white arrows) intermixed with occasional mouse nuclei (punctate, condensed chromatin) as further confirmation of human derived cells engrafted into the murine epithelial microenvironment. (**I-J**) Engrafted gcCFTR UABCs from second CF donor 1444, with expression of both human nuclear specific marker Ku80 (white) and human specific EpCAM (EpCAM Hu, green), with mouse specific EpCAM again confined to the niche periphery. (J) Differentiated human secretory cells (MUC5AC, green) and ciliated cells (α-tubulin, red) are detected within the niche, along with human specific nuclear marker Ku80 (white), co-stain in engrafted regions of CF donor 1444. (**K**) Larger human derived gcCFTR UABC cells also show abundant expression of MUC5AC (arrows) compared to endogenous mouse derived goblet cells (arrow heads). A dramatic difference in cell dimensions can be noted between human and endogenous mouse goblet cells (arrows, 24.17 +/- 1.26). (**L-M**) Non-CF donor 1453 show similar engraftment and staining patterns to CF donors. (**L**) Truncated CD19 (tCD19, green) human cells are flanked by mouse specific EpCAM (EpCAM Ms, red) expression. (**M**) Pronounced cell hypertrophy from presence of mucin (MUC5AC, green) in non-CF human gcUABCs is similar to CF donor gcUABCs following engraftment (H, K) while ciliated cells appear unchanged (α-tubulin, red). Scale bars: (E-G) 25 µm, (H and K)10 µm, (I) 20µm (J, L, M) 25 µm.

The nasal epithelium of immunocompromised NSG mice was preconditioned using PDOC, and gcUABCs from CF and non-CF donors were transplanted into the upper airways and tracked over several months. To distinguish between non-fluorescent human and murine cells in the chimeric nasal epithelial tissues in transplanted mice, sequential serial sections of transplanted mice were immunohistochemically assayed using species specific expression of epithelial cell adhesion molecule (EpCAM) in conjunction with antibodies against CD19. Figure 4E-H shows gcUABCs from one representative CF donor 2.5 months following xeno-transplantation. Murine specific EpCAM sharply discriminated the murine epithelium and demonstrated niches devoid of staining. Human specific EpCAM overlapping with tCD19 confirmed these niches as focal sites harboring human gcUABCs (Fig. 4E, F), confirming successful human stem cell engraftment by immunohistology. Differentiated progeny human ciliated and secretory cells were additionally detected 75 days-post-transplantation (Fig. 4G). Human nuclei from engrafted airway stem cells were additionally distinguished from recipient murine nuclei given absence of condensed nuclear chromatin plaques in the former (Figure 4H (*37*)). Interestingly, transplanted human gcUABCs assumed the typical multilayered, pseudostratified architecture seen in normal human sinonasal tissue (Fig. 4G, H), and prominent expression of MUC5AC in engrafted cells (Fig. 4H). Transplanted gcUABCs from a second CF donor demonstrates a niche of engrafted cells with colocalized staining of human specific EpCAM and human nuclear protein Ku80 (Fig. 4I) Staining of murine specific EpCAM is restricted to the perimeter of chimeric niches following PDOC preconditioning. Pronounced secretory cell hypertrophy (∼24x enlarged compared to resident murine cells) was again observed (Fig. 4J, K). Control human non-CF gcUABCs showed identical patterns of biomarker staining within the engrafted niches as CF donors (Fig. 4L), with prominent secretory cell hypertrophy and MUC5AC expression again seen (Fig. 4M).

### Engrafted human gcUABCs maintain expression of engineered CFTR and CD19 proteins *in vivo*

We assessed the expression of engineered CFTR and CD19 proteins following human gcUABC transplantation into the recipient murine background. Anti-human CFTR antibody ACL-006 was selected for downstream experiments given robust, specific staining of CFTR protein in normal mouse lung (Fig. 5A), and absence of staining in lungs of CFTR knockout mice (Fig. 5B). When focusing solely on the engrafted niches in recipient mice that harbor human gcUABCs, the expected multi-tiered normal architecture of human epithelium is again reconstituted, as is expression of Krt5 and engineered tCD19 transmembrane protein at the cell surface (Fig. 5C). Maintained expression of engineered wild-type CFTR protein is detected in niches with a high proportion of mucin-producing cells (Fig. 5D), or is observed in a punctate pattern within non-ciliated cells, with minimal transgene expression in occasional differentiated human ciliated cells (Fig. 5E). These data collectively suggest the retention of native stem cell properties of human airway stem cells (such as *in vivo* epithelial organization and differentiation) following both *CFTR* gene correction and long-term engraftment into the chimeric murine microenvironment. The differentiated cell progeny of human stem cells also maintain the robust expression of the desired engineered proteins *in vivo,* uninfluenced by the immunosuppressed, xenogeneic background with no aberrant cell transformation observed.

**Fig. 5:**
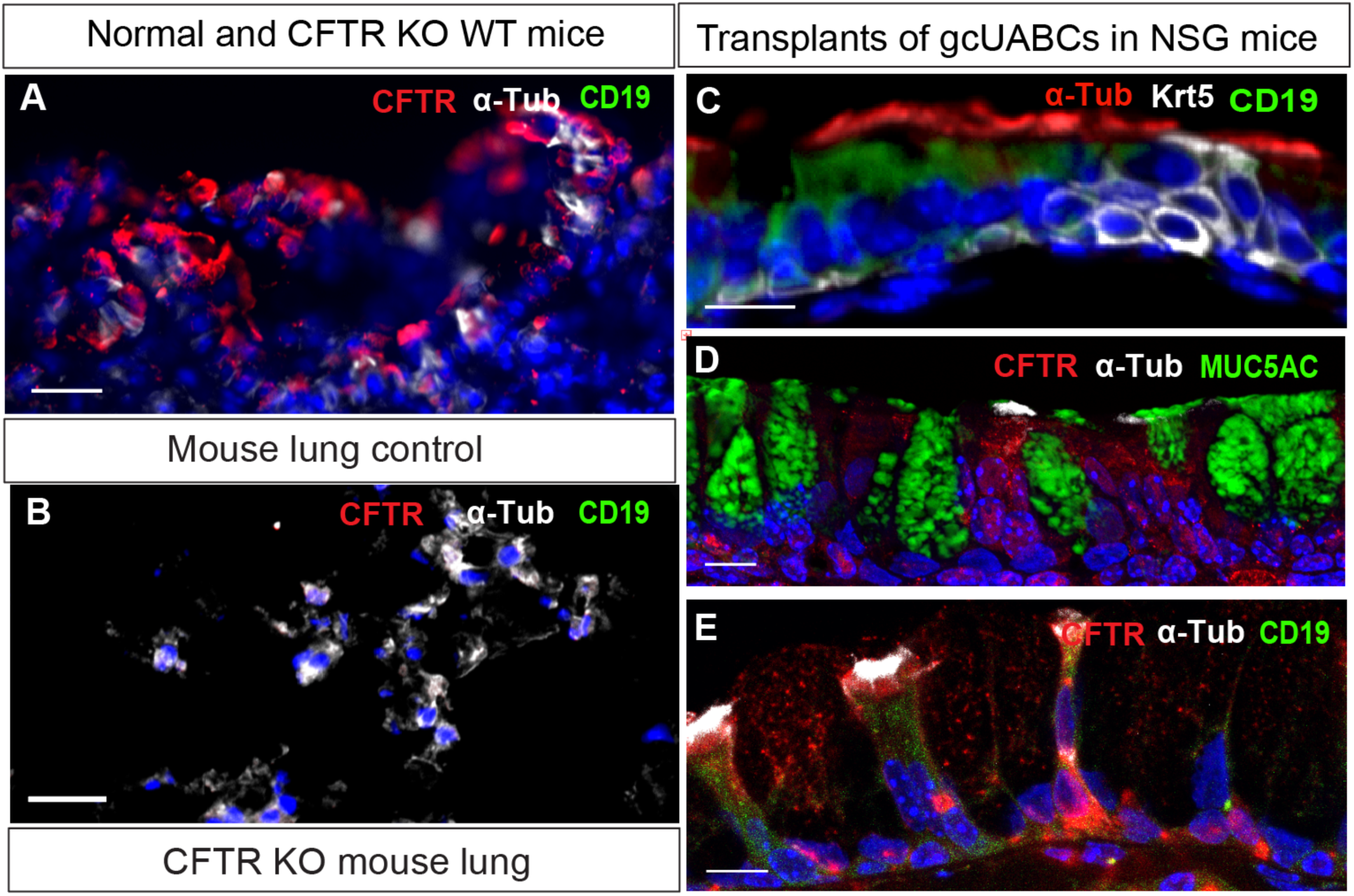
Retained expression of CFTR and CD19 proteins following gcUABC chimeric engraftment. (**A, B**) Validation of CFTR protein detection conferred using anti-CFTR antibody (ACL-006) to discriminate CFTR expression (red) in wild-type murine lung (A), which is absent in CFTR knock-out murine lung (B). (**C**) Following transplant of gcUABCs from CF donor 1444, CD19 (green) and α-tubulin (red) expression were both detected at low magnification. Multi-tiered Krt5 stem cells (white), characteristic of normal architecture in human sinonasal mucosa, are noted following cross-species engraftment. (**D**) A broad fluorescence pattern of CFTR expression (red) is detected from the hypertrophied secretory cells (MUC5AC, green) derived from gcUABCs. (**E**) In the absence of MUC5AC, punctate CFTR expression is readily detected by goblet cells (red) along with cell surface CD19 expression (green). CFTR staining is noted in the niches from gcUABCs stem cell engraftment. This was primarily seen in apical, engorged secretory cells as well as some gene edited ciliated cells (α-tub, white). Scale bar: 10 µm (A-E).

## Discussion

Given the substantial safety hurdles inherent to the delivery and engraftment of airway stem cells into the lungs of CF patients, we have advanced the concept of *ex vivo CFTR* gene correction in primary UABCs sampled from CF donors, followed by autologous, transplantation of gene corrected, functional UABCs into donor upper airways. Unlike the sinus epithelium which can be safely denuded, substantial acute debridement of the lower airways could result in respiratory insufficiency. Several CF gene therapy trials using viral or non-viral vectors to deliver CFTR cDNA into resident cells in the upper (nose and sinus) and lower airways have been unsuccessful (*19*, *21*, *38*), likely due to poor stem cell transduction and limitations after repeat treatment dosing from acquired viral vector immunity (*22*, *38*). *In vivo* immune responses against gene-editing enzymes such as CRISPR-associated protein 9 (Cas9) *in vivo* limit the persistence of gene-corrected cells in murine (*24*) and canine models (*39*). These experiences advocate for an approach towards stem cell gene correction achieved ex-vivo (*27*), followed by autologous stem cell transplantation and engraftment into sites of active regional disease (*23*, *39–41*). We have focused efforts towards the treatment of CF-CRS, since primary UABC stem cells are abundant and accessible for gene correction, and the sinuses as targets for cell delivery are tolerant to tissue manipulation and post-transplant surveillance without compromising patient safety.

CF patients are highly impacted by recurrent CF-related chronic rhinosinusitis (CF-CRS), typified by nasal inflammation, impaired mucociliary clearance, abundant intranasal scarring, and recurrent bacterial abscesses within the sinuses (*42*, *43*). Furthermore, the diseased upper airway and sinuses harbor pathogens that are drug resistant and seed subsequent infections of the lungs and/or transplanted lungs, thereby accelerating lower airway deterioration (*44*, *45*) (*46*). Restoration of CFTR function to the sinus epithelium using combined genetic and stem cell therapies holds promise in mitigating CF-CRS in patients with CF who have a contraindication or intolerance to modulator use and may sustain and improve lung health either prior to or following lung transplantation in modulator-intolerant patients.

We have previously reported our large transgene insertion strategy for *ex vivo* CFTR restoration to insert wild-type CFTR cDNA using CRISPR-Cas9 RNP and two simultaneously delivered adeno-associated viruses (AAVs) to upper and lower airway basal stem cells. This methodology is agnostic to specific CFTR mutations and should theoretically be applicable to virtually any CF-causing mutation, including unidentified and class I CF mutations. We directly insert the full-length CFTR cDNA at the start codon in *CFTR* exon 1 using Cas9 and AAV to restore CFTR function to physiological levels observed in non-CF controls (*27*). This ‘super-exon’ approach to place wild-type cDNA into the endogenous CFTR start codon rescues all downstream disease-causing mutations while preserving both endogenous regulation of the gene and overall genomic structure (*47*). Additionally, we sequentially insert a truncated CD19 (tCD19) enrichment tag into human sinonasal UABCs (*27*), which has no physiologic signaling because the intracellular domain is removed. tCD19 is used here to both track the knock-in cells with immunofluorescence microscopy, and also enrich for knock-in UABCs using magnetic beads in which 60-80% enrichment was achieved with variance between disparate CF donors harboring a wide spectrum of CFTR mutations. Restored physiologic CFTR chloride transport function in the mature airway progeny cells of gcUABCs was found to be similar to anion transport seen from donors without CF in ALI cultures, exceeding predicted efficiencies that would be required for clinical scale application (*27*).

Regarding *in vivo* debridement for these studies, the use of detergent (2% polidocanol) for cell transplant was recently reported as showing limited success (*32*). However, those experiments were performed without the use of a scaffold or matrix material to adhere the transplanted cells *in situ*. In our early work, we observed the leakage of transplanted cells from the mouse nasal cavity. Here, we show our efforts to optimize injury methods and screen for clinically applicable scaffold biomaterials to achieve reliable transplantation of mouse and human upper airway basal stem cells (UABCs) in immunocompromised mice NOD *scid* gamma (NSG^TM^) mice. We prioritized substrates conducive for the proliferation of UABCs and stem cell maintenance, or clinically compatible materials reported in tissue engineering applications. Among them, the optimal substrates of fibrinogen/thrombin for UABCs engraftment are noteworthy as they are native human proteins involved in clot formation and are used in sinus surgical procedures.

Another consideration in the development of this stem cell therapy is the percent of the epithelium that must be replaced. In our previous *in vitro* studies, we observed robust restoration of CFTR function to 30-60% of the levels in non-CF controls when only >20% of cells were corrected (*27*). Although we obtain >50% corrected UABCs *in vitro*, the percent of cells that successfully engraft in this pre-clinical, xenogeneic system may be lower. The mouse-to-mouse transplantation experiments show that about 30% of the epithelium can be replaced in mice following topical pre-conditioning. However, the diminutive nasal passages in mice prevented surgical placement or implantation of the UABCs encapsulated in fibrinogen to the site of injury. Since future anticipated procedures in candidate CF patients would be performed using endoscopic equipment, directed debridement of niches with targeted delivery of gcUABCs to the precise site undergoing conditioning are expected to improve the efficiency of transplantation.

Apart from the efficacy of this approach, the safety profile of genome-edited UABC transplantations should be emphasized. Our *in vitro* studies have to date been devoid of enrichment of UABCs with oncogenic mutations (*27*), and we have not observed any tumor formation in mice engrafted with genome edited human UABCs. Future preclinical studies will use clinical grade cell therapy products to maintain safety during transplantation of genome edited human UABCs.

The secretory cell hypertrophy seen in human mucin 5AC differentiated cells following xenogeneic engraftment is seen in both gene corrected CF and non-CF human donor samples. This degree of MUC5AC expression may be related to variability of transplantation between species, and/or murine microenvironment interactions. Notably, *in vitro* primary CF UABCs will differentiate into a predominantly mucosecretory tissue, as observed here (48). These observations suggest that the proliferative and differentiation program of the UABCs is preserved and not altered upon editing and subsequent transplantation.

The limitations of this study include that while expression of human CFTR was detected in the correct cells, CFTR function could not be assessed following *in vivo* engraftment because the NSG mice express native wild-type CFTR. Additionally, this study was performed in mice with genetically-conferred immunomodulation. Before trial in CF patients, this protocol would ideally be validated in an additional model, with sinuses of a scale compatible to humans, although the necessary degree of sufficient immunomodulation to prevent rejection of transplanted human cells would need to be determined. It is also conceivable that the local presence of sinonasal inflammation may limit the engraftment of our gene corrected UABCs in a clinical trial of people with CF. However, the anticipated surgical debridement process in humans would remove infected and inflamed sinus tissue to mitigate these concerns.

The results presented here validate a graduated approach to achieve sustained engraftment of exogenously transplanted mouse and human UABCs into the airway epithelium *in vivo*. Moreover, the gene corrected human CF UABCs maintain the ability to differentiate into the major ciliated and secretory cells of the normal epithelium, and faithfully express CFTR protein from our spliced-in transgene. Our findings thereby demonstrate the restoration of CFTR function in respiratory epithelium derived from gene corrected UABCs *in vitro* and proof of engraftment of these gene-corrected UABCs *in vivo*. Of note, evidence of CFTR restoration *in vitro*, along with proof of safety *in vivo*, was sufficient to proceed with clinical trials for CFTR modulators. Thus, our studies present a rational and promising pre-clinical springboard for testing of autologous gene corrected airway stem cell therapies to treat CF-CRS in clinical trial. If successful, the knowledge and expertise developed in the implementation of an upper airway stem cell transplantation therapy might also be adapted towards CF lower airway applications.

## Supporting information

Suppl. Bravo et al.

## Acknowledgments

The authors would like to thank Manasvi Marathe, Rhonda Perriman, and Helen Simon for their organizational and grant writing efforts with this collaborative group as well as assistance with assembly of this manuscript.

## Funding that supported this work

National Heart, Lung and Blood Institute grants R01 HL151677 to JVN; 1K99HL151900-01A1 to SV. Cystic Fibrosis Foundation grants and Ross Mosier Research Laboratories gift fund to CM. Cystic Fibrosis Research Institute grant to MHP.

## Author contributions

Conceptualization: DTB, SV, JB, RSN, JJW, CM, ZMS, TJD, MHP, JVN

Methodology: DTB, SV, JB, VS, ET, PR, WWK, PJA, ZMP, PHH, RSN

Investigation: DTB, SV, JB, VS, ET, PR, WWK, PJA, ZMP, PHH, RSN

Visualization: SFB, MJM, JLS, EH

Funding acquisition: DTB, SV, TJD, MHP, JVN

Project administration: TJD, MHP, JVN

Supervision: DTB, SV, JB, RSN, TJD, MHP, JVN

Writing – original draft: DTB, SV, JB, VKR, ZMS, TJD, MHP, JVN

Writing – review & editing: DTB, SV, VKR, JJW, CM, ZMS, TJD, MHP, JVN

**Supplemental Fig. S1: Debridement methods demonstrate epithelial disruption.** (**A**) Control mice prior to SO_2_ gas exposure. (**B**) Mice exposed to SO_2_ gas for 5 hours. Epithelial disruption is observed 24 hours post exposure. (**C**) 7 days post injury, epithelial regeneration was apparent. (**D**) Control mouse prior to PDOC exposure. (**E**) To promote engraftment of transplanted UABCs, 2% polidocanol injury caused epithelial disruption within 24 hours post injury. (**F**) Natural epithelial repair/regeneration 7 days post injury in the upper airway. Scale bar: 100µm.

**Supplemental Fig. S2: Engraftment of transplanted GFP/Luc UABCs following chemical debridement.** (**A**) Transplantation of GFP/Luc UABCs after SO_2_ gas exposure for 5 hours at 6- and 19-hours post injury. BLI in radiance (photons/sec) was measured over 6 months. (**B**) Stable engraftment was observed compared to controls (p < 0.0001). (**C**) Immunofluorescence of an SO_2_ treated mouse transplanted with GFP/Luc UABCs. Scale bar: 50 µm.

**Supplemental Fig. S3: Long term engraftment of GFP/Luc+ UABCs after mechanical preconditioning of the upper airways.** (**A**). Stable engraftment of UABCs derived from transgenic reporter mice constitutively expressing GFP/Luc to the upper airways of NSG mice following preconditioning by nasal brush (NB) application. BLIs demonstrate stable and durable engraftment for over 6 months as compared to control (p > 0.0001). (**B**) Mechanical debridement by NB, achieved epithelial disruption to enable lasting engraftment of GFP/Luc UABCs to the upper airways as demonstrated by immunofluorescence. GFP/Luc UABCs became incorporated into the basement membrane as a basal stem cell monolayer (Krt 5, white) and progeny differentiated cells expressing GFP produced an overlying pseudostratified epithelium. Isolated GFP^+^ basal cells are also seen within the epithelium (arrows). (**C**) Magnified image of dashed box in (B). As GFP expressing cells are not continuous but the BLI measurements are stable, UABCs engraft to a particular location and self-renew, as opposed to spreading throughout the epithelium. Scale bar: 100 µm (B), 25µm (C).

**Supplemental Fig. S4: Transplantation of human gcUABCs into immunocompromised mice.** UABCs from donors were edited to express Nanoluc^TM^ luciferase and GFP and then transplanted into several preconditioned NSG mice. Transplanted mice exhibited stable bioluminescence for >60 days.

## Materials and Methods

### Culture and genome editing of UABCs

GFP-Luciferase (GFP-Luc) UABCs were isolated from the nasal cavities of L2G85 transgenic mice (*28*, *29*). Human sinonasal tissue from people with and without CF was obtained following functional endoscopic sinus surgeries (FESS) using approved institutional review board protocols. The process for culturing and genome editing UABCs has been previously described by our group (*26*). Briefly, whole sinonasal tissues were suspended in pronase (1.5 mg/mL)/F-12 media for 2 hours at room temperature or overnight at 4°C. After passage through a cell strainer, total cells were resuspended in Pneumacult X-plus medium with ROCK inhibitor, Y-27632 at 5% O_2_ on tissue culture plates coated with recombinant laminin-511 (iMatrix^TM^). Enriched, adherent UABCs were expanded 5 days after primary isolation. A single guide RNA (5’-CTTGCCCCACAGGGCAGTAA-3’) specific to the hemoglobin B (HBB) locus not expressed in this cell type or lineage (‘safe harbor’ for insertion) was used to induce a double stranded break (DSB). In selected experiments, to track transplant efficacy and cell localization, adeno-associated virus (AAV) carrying a DNA template encoding for both Nanoluc^TM^ and TurboGFP under the control of the spleen focus forming virus (SFFV) promoter was used to edit human UABCs. Edited UABCs were passaged 7-10 days, and GFP+ UABCs were enriched using fluorescence-activated cell sorting. In experiments inserting the CFTR cDNA, double strand breaks (DSBs) were induced by a single guide RNA (sgRNA) specific to exon 1 of the CFTR locus (5’-TTCCAGAGGCGACCTCTGCA-3’). We have previously described use of two AAVs to sequentially insert two halves of the full length CFTR cDNA plus a truncated CD19 (tCD19) tag under a PGK promoter (***26***). Edited UABCs expressing tCD19 were instead enriched using magnetic activated cell sorting (MACS). Cells were cultured for 3-5 days after cell enrichment before transplantation.

### Testing of candidate scaffold material

Human fibrinogen (Sigma-Aldrich) was freshly dissolved in phosphate-buffered saline (PBS) at a concentration of 3 mg/mL prior to each in vitro and in vivo experiment. Fibrinogen precipitates from the solution if left overnight. Thrombin (Sigma-Aldrich) was dissolved in water at a concentration of 2 units/mL. Thrombin was aliquoted and frozen at −20°C for long-term use. Fibrinogen was added to a cell pellet and cells were resuspended. Thrombin was added immediately prior to plating or transplantation. For in vitro experiments, the final volume of the gel was 40 mL. For in vivo experiments, 15 mL of fibrinogen plus thrombin mixed with cells was used. Table 1 lists other scaffold materials with usage advantages and concentrations examined.

### Pre-conditioning and transplantation procedures

All animal procedures were approved by the Animal Care and Use Committee (IACUC) at Stanford University. Mice were obtained from Jackson Labs (Bar Harbor, ME USA) and used at 8-12 weeks old. Immunocompromised NOD *scid* gamma mice (NSG) mice were housed in the Stanford University animal care facility. For nasal brushing mechanical preconditioning techniques, mice were anesthetized using ketamine (80–100 mg/kg) and xylazine (8–10 mg/kg). Interdental brushes (0.5 mm diameter) were used to disrupt the epithelial lining of the cavity via gentle twist insertion of the brush to a depth of about 8-10 mm. Two hours post-conditioning, UABCs embedded in Matrigel^TM^, in a 10 μL volume were added unilaterally to the upper airway cavity. Chemical preconditioning of the upper airway epithelium was performed using SO_2_ at 500 ppm balanced air (ProSpec^TM^, Praxair) in a 2-liter induction chamber. Flow rate was held at approximately 4.3 standard cubic foot per hour for 5 hours. After 6 or 19 hours, ∼ 0.5-1×10^6^ UABCs embedded in Matrigel^TM^, were added unilaterally to the upper airway cavity. Preconditioning with 2% polidocanol (PDOC), was performed on anesthetized mice, intranasally to a single side of nasal cavity to promote engraftment to the respiratory epithelium. NSG mice expressing human IL3, GM-CSF (CSF2) and SCF (KITLG) (NSG-SGM3) were used. The mice were allowed to recover for 2 hours post treatment. UABCs were embedded with the appropriate biomaterial and 10-15 mL was intranasally administered into the conditioned airway cavity. Mice were monitored for improper breathing. Bioluminescence was monitored at reported intervals as described below.

### In vivo bioluminescence imaging (BLI)

Studies evaluating the transplantation utilized UABCs from mice expressing firefly luciferase. Firefly luciferase activity was measured after the injection of D-luciferin intraperitoneally. Human UABCs were edited to express Nanoluc^TM^ and TurboGFP. Nanoluc activity was measured using the substrate Nanoglo^TM^ (fluorofumarizine). One vial of Nanoglo was resuspended in 525 mL PBS and 50 mL was injected intraperitonially per mouse. Mice were recovered from anesthesia for 5 minutes, then re-anesthetized using isoflurane to measure bioluminescence using a Lago imager. Images were obtained with a 300s exposure.

### Tissue processing and immunohistochemistry

The skulls/upper airway cavities from NSG mice, at selected time points, were fixed in 4% paraformaldehyde (Electron Microscopy Services, Hatfield, PA) in PBS (pH 7.4) at 4°C overnight. Samples were treated on alternating days with 0.5 mM EDTA in PBS at 4°C as required to achieve decalcification. Tissues were cryoprotected by successive incubations in 15%, 20%, and 30% sucrose in PBS. Samples were embedded in 100% OCT compound (Sakura Finetek, Torrance, CA) following 30 min successive incubations in 30%, 50%, 90% OCT in 30% sucrose/PBS at room temperature. Serial frozen sections of 10 μm thickness were obtained using Leica cryostats CM1950/3050S (Leica, Bannockburn, IL). Non-specific binding sites were blocked in PBT1 consisting of 0.2% Triton X-100 (EMD Chemical Inc., Gibbstown NJ), 1% bovine serum albumin (Fisher Scientific, Fair Lawn, NJ), and 0.02% NaN_3_ (EMD Chemical Inc., Gibbstown NJ) in PBS, for 1 hour at room temperature. The tissues were then treated overnight at 4°C in PBT2 buffer containing of 0.2% Triton X-100, 0.5% bovine serum albumin and 0.02% NaN_3_ in PBS with the following primary antibodies: acetylated-α-tubulin (1:200, Cell Signaling, Danvers, MA; Millipore-Sigma Temecula, CA), MUC5AC (1:200), Ki67 (1:200), and GFP (1:500), Abcam, Cambridge, MA), anti-mouse EpCAM, anti-human EpCAM anti-human CD19 (1:200) (BD Biosciences, San Jose, CA), Ku80 (1:200, Bioss, Woburn, MA), Turbo-GFP (Invitrogen Corp., Carlsbad, CA) and Krt5 (BioCare Medical LLC, Pacheco, CA). The following day, tissues were washed 3 times for 10 minutes with PBS. Tissues were then incubated with secondary antibodies, Alexa Fluor 488, 546, and 647 (1:500, Molecular Probes, Invitrogen Corp., Carlsbad, CA) or Alexa Fluor 488 1:500, Jackson ImmunoResearch, West Grove, PA) in PBT3 buffer containing 0.2% Triton X-100, and 0.02% NaN_3_ in PBS for 1-2 hours at room temperature. Nuclei were stained with Hoechst in PBS (1:2000, Invitrogen Corp., Carlsbad, CA). Tissues were then mounted using antifade fluorescent mounting media (DAKO, Carpinteria, CA). Tissue sections were imaged using a Zeiss Axioimager/LSM 5 Exciter fluorescence microscope, and a Carl Zeiss LSM880 confocal microscope.

For IHC protein staining of histo-sections of xeno-transplanted and engrafted human gcUABCs on mouse backgrounds, the following anti-human CFTR antibodies and concentrations were tested for staining efficacy: i) CF Foundation, Chapel Hill, NC, CFTR distribution program, (1:100). ii) CF Foundation, Chapel Hill, NC, (1:100). A6-450. iii) Alomone Labs, Jerusalem, IL, ACL-006 (1:100). Ultimately, the clone from Alomone Labs was used for staining in Figure 5 as described.

### Engraftment analysis

To determine the percent engraftment of transplanted murine UABCs, the final GFP incorporation was determined on the day of sacrifice. This time point for murine GFP+-Luc+ cell transplants, was from 7-9 months post-transplant. Each mouse upper airway cavity was serially sectioned. Images were stitched into a composite view of the entire cavity. Using image J software, the length of the cavity was determined by tracing the epithelial surface. The percent engraftment was determined by the length of GFP expressing cells (cumulatively) divided by the length of the cavity, minus the nasal floor and the olfactory areas (devoid of engrafted cells). This was performed for each mouse of the experiment. After performing this analysis, efficiency of cell number transplanted in male and female mice was calculated by the ratio of the length of GFP expression (measured cumulatively, in a 10µm section), to the length of the nasal cavity, between the OE and NF (respiratory epithelium).

