## Supplementary material for "Durable reconstitution of sinonasal epithelium by transplant of CFTR gene corrected airway stem cells": Suppl. Bravo et al.

**Supplemental Figures S1-S4:**

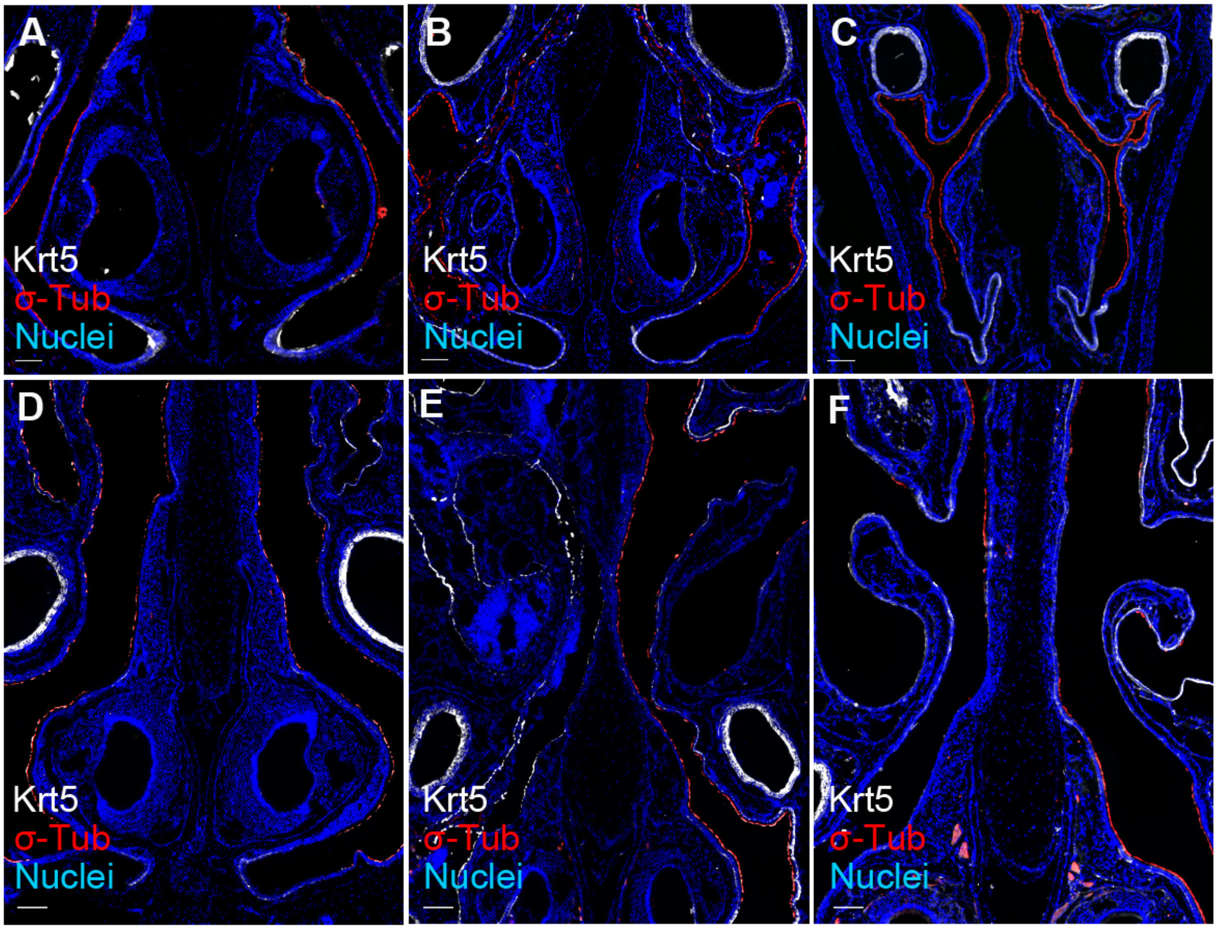


**Fig. S1. Debridement methods demonstrate epithelial disruption.**

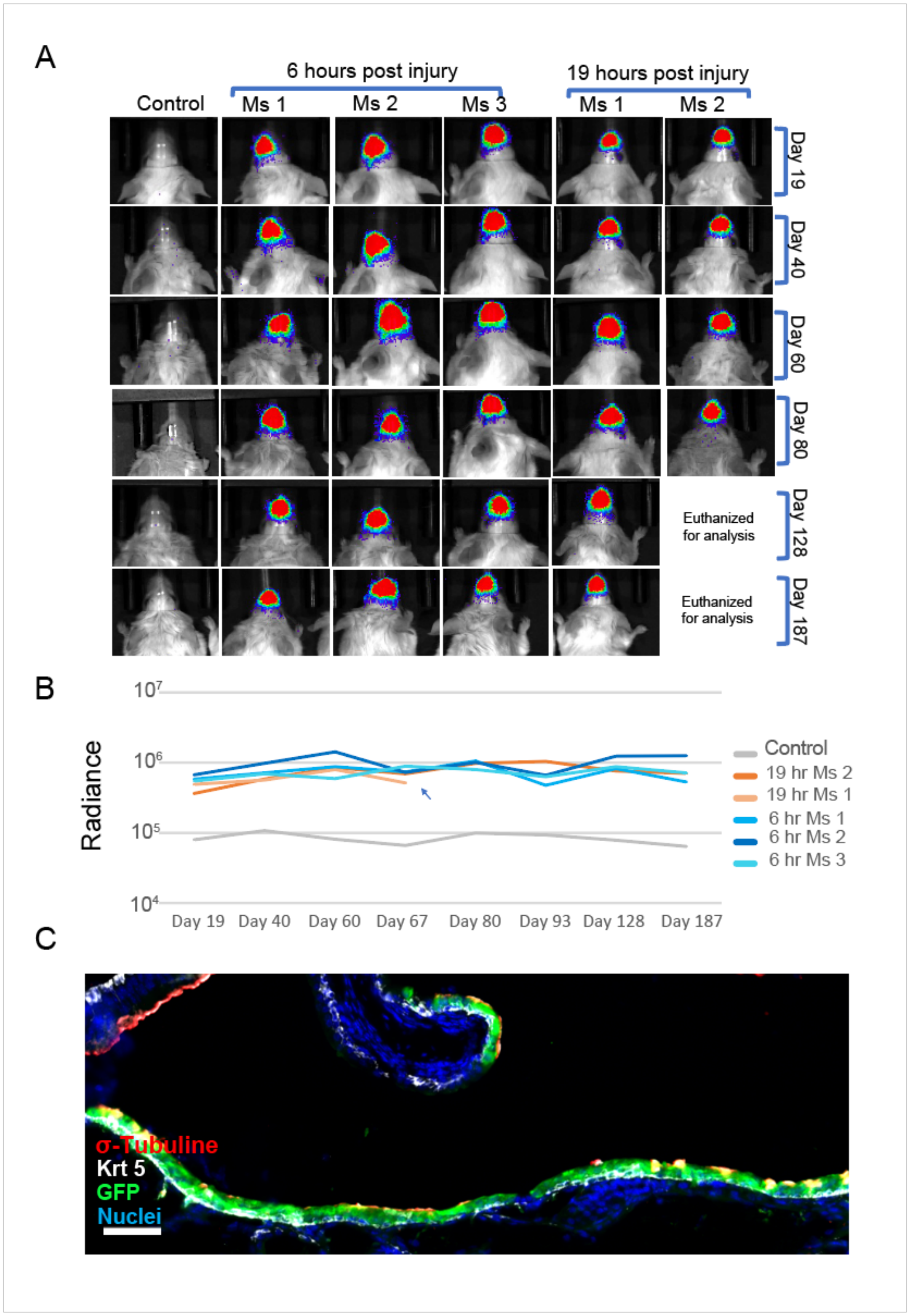


**Fig. S2. Engraftment of transplanted GFP+/Luc+ UABCs following chemical debridement.**

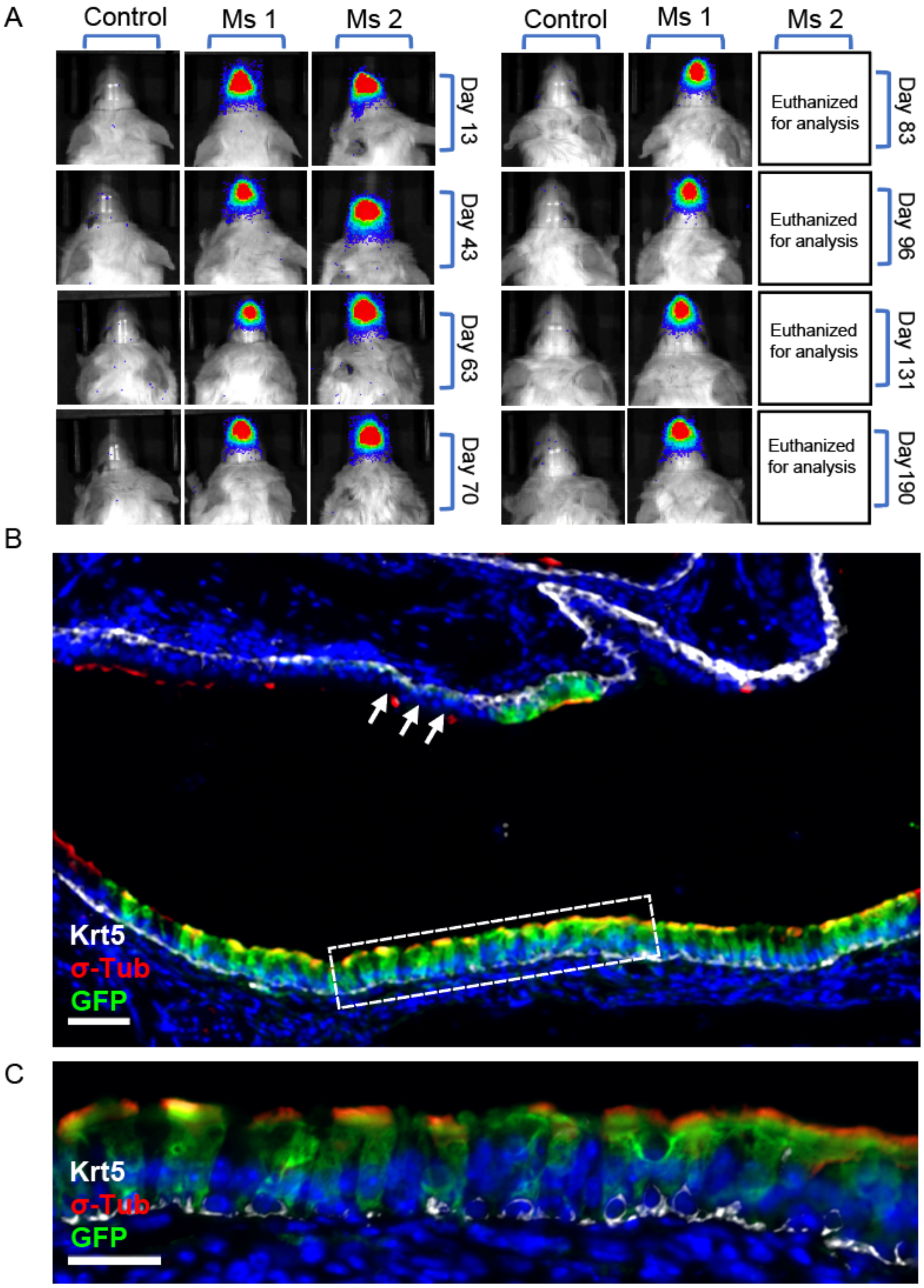


**Fig. S3. Long term engraftment of GFP+/Luc+ UABCs after mechanical preconditioning of the upper airways.**

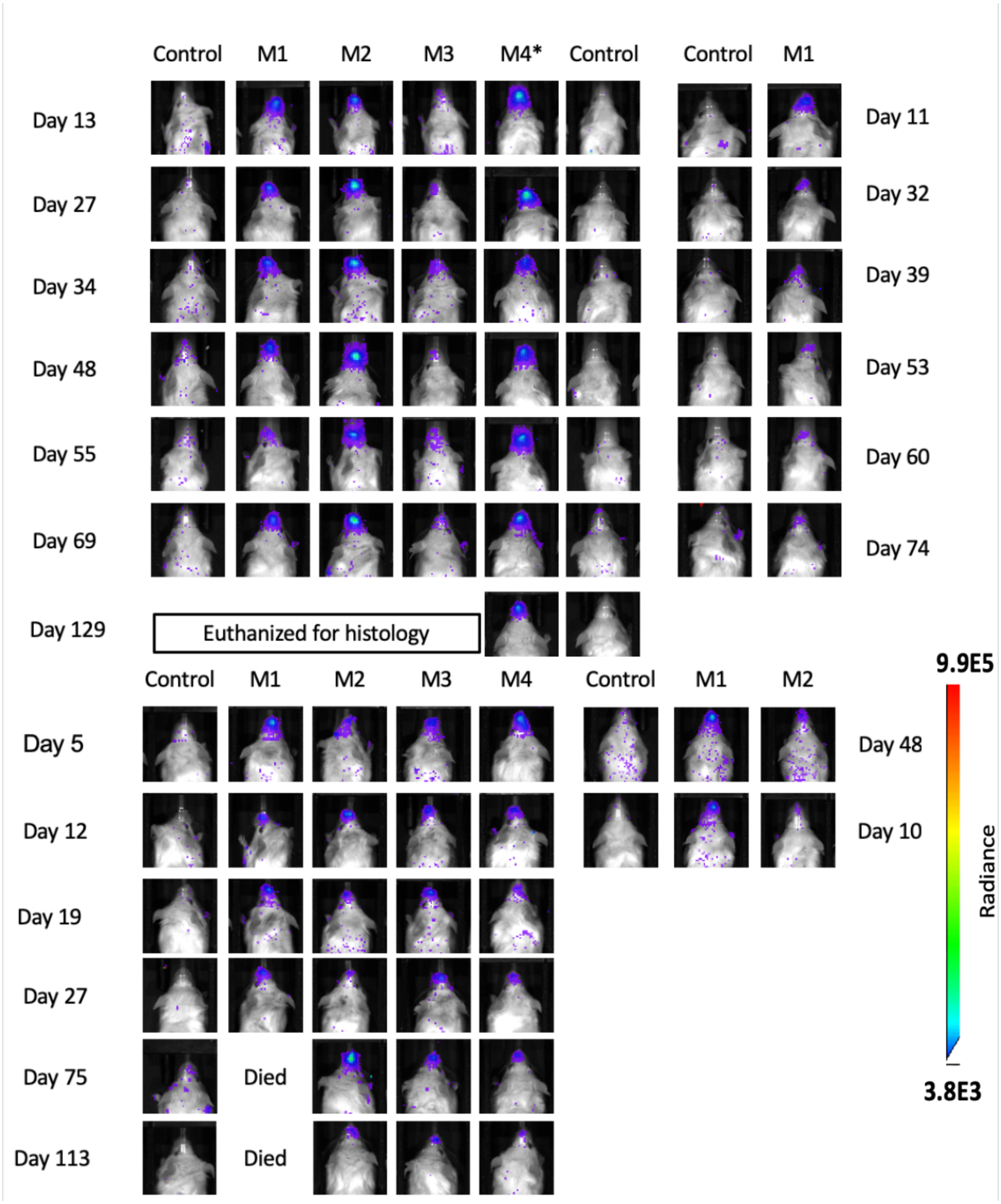


**Fig. S4: Transplantation of human gcUABCs into immunocompromised mice.**

| Sample No. | Material | Advantages | Optimal concentration |
|---|---|---|---|
| 1 | Fibrinogen | Human protein produced in the liver Component in common surgical biomaterials/adhesives | 2.85 ug/μL |
| 2 | Spidersilk functionalized with laminin (BioSilk™) | Laminin coated matrix for cell adhesion and proliferation | 3 ug/μL |
| 3 | Bovine Type I collagen (PureCol™) | Extracellular matrix material with widespread clinical use | 3 ug/μL |
| 4 | Hyaluronan (HyStem-C™) | Biocompatible polymer implicated in epithelial mucosal repair | 5 mg/mL |
| 5 | RGD functionalized Dextran (TrueGel™) | Biocompatible polymer used to encapsulate cells for growth | Proprietary |
| 6 | Alginate | Inert plant-based polymer currently used in food products | 20 mg/mL |

**Table S1. Biomaterials evaluated for human UABC transplant**

| Donor | Donor ID/Exp ID | Genotype | tCD19+ cells | CFTRinh-172 response |
|---|---|---|---|---|
| 1 | CF1413/SV390 | F508/F508 | 61% | 9.6 |
| 1 | CF1413/SV390 | F508/F508 | 61% | 4.3 |
| 2 | CF1418/SV391 | Exon14 deletion homozygous | 57% | 3.06 |
| 2 | CF1418/SV391 | Exon14 deletion homozygous | 57% | 2.92 |
| 3 | CF1444/SV405 | F508del/3659delC | 57% | 28 |
| 3 | CF1444/SV405 | F508del/3659delC | 57% | 27 |
| 3 | CF1444/SV405 | F508del/3659delC | 57% | 22 |
| 4 | CF1471/SV416 | dF508/W1204X | 53% | 4.6 |
| 4 | CF1471/SV416 | dF508/W1204X | 53% | **3.0** |

**Table S2. gcUABC CF Donors.**

Summary of CF donor genotype, % allelic correction and CFTR microcurrent responses in epithelial air liquid interface cultures *in vitro* following gene correction of human UABCs
